# Metatranscriptomic comparison of endophytic and pathogenic *Fusarium*–Arabidopsis interactions reveals plant transcriptional plasticity

**DOI:** 10.1101/2021.03.01.433437

**Authors:** Li Guo, Houlin Yu, Bo Wang, Kathryn Vescio, Gregory A. DeIulio, He Yang, Andrew Berg, Lili Zhang, Véronique Edel-Hermann, Christian Steinberg, H. Corby Kistler, Li-Jun Ma

## Abstract

Plants are continuously exposed to beneficial and pathogenic microbes, but how plants recognize and respond to friends versus foes remains poorly understood. Here, we compared the molecular response of *Arabidopsis thaliana* independently challenged with a *Fusarium oxysporum* endophyte Fo47 versus a pathogen Fo5176. These two *Fusarium oxysporum* strains share a core genome of about 46 Mb, in addition to unique 1,229 and 5,415 accessory genes. Metatranscriptomic data reveal a shared pattern of expression for most plant genes (∼80%) in responding to both fungal inoculums at all time points from 12 to 96 h post inoculation (HPI). However, the distinct responding genes depict transcriptional plasticity, as the pathogenic interaction activates plant stress responses and suppresses plant growth/development related functions, while the endophytic interaction attenuates host immunity but activates plant nitrogen assimilation. The differences in reprogramming of the plant transcriptome are most obvious in 12 HPI, the earliest time point sampled and are linked to accessory genes in both fungal genomes. Collectively, our results indicate that the *A. thaliana* and *F. oxysporum* interaction displays both transcriptome conservation and plasticity in the early stages of infection, providing insights into the fine-tuning of gene regulation underlying plant differential responses to fungal endophytes and pathogens.

**One-sentence summary:** Multiomics analysis reveals the regulatory plasticity of plants in response to beneficial and antagonistic microbes, resulting in distinct phenotypes and rewired transcriptional networks.

## INTRODUCTION

Over millions of years of coevolution, plants and microbes have established intimate relationships, forming beneficial, neutral, or antagonistic partnerships. Plant pathogens threaten agricultural production and global food security (Dean et al., 2012; Strange and Scott, 2005), but beneficial microbes, such as rhizobia, mycorrhizae, and endophytes limit plant pests and promote plant growth through nutrient mineralization and availability (Rashid et al., 2016; White et al., 2019). How plants recognize and react differently to friends versus foes is an intriguing topic of research.

Our understanding of plant immunity has been revolutionized by the recent increase in the breadth of genomic data available. The classical, binary view of plant immunity consists of pattern-triggered immunity (PTI) and effector-triggered immunity (ETI) (Jones and Dangl, 2006; Cui et al., 2015). Plant PTI relies on plasma membrane (PM)-localized pattern recognition receptors (PRRs), which are often receptor-like proteins/kinases (RLPs/RLKs) that sense conserved microbe-associated molecular patterns (MAMPs) or damage-associated molecular patterns (DAMPs) and induce downstream defense reactions (Bigeard et al., 2015; Dodds and Rathjen, 2010; Jones and Dangl, 2006; Zhou et al., 2020). Plant ETI employs an intracellular nucleotide-binding site and leucine-rich repeat domain receptors (NLRs) that recognize specific microbial effectors as a result of ongoing host-pathogen coevolution (Jones and Dangl, 2006; Cui et al., 2015; Wang et al., 2020b; Cesari, 2018; Monteiro and Nishimura, 2018). The oftentimes blurry boundary between MAMPs and effectors prompted a new spatial immunity model based on extracellular or intracellular locations of pattern recognition (Wang et al., 2020b; van der Burgh and Joosten, 2019).

The comparative study by Baetsen-Young and colleagues (2020) revealed transcriptional reprogramming of *Fusarium virguliforme* when interacting with different plant hosts (soybean versus maize). This study investigated transcriptome reprogramming of the same plant host and explored how PTI and ETI or extracellular and intracellular immunity are involved in both beneficial and antagonistic interactions. We established the *Fusarium oxysporum*–*Arabidopsis thaliana* model system, which includes an endophyte, *F. oxysporum* strain Fo47, and a pathogen, *F. oxysporum* strain Fo5176. Arabidopsis plants infected by these two *F. oxysporum* strains display distinctive phenotypes, with Fo5176 causing typical vascular wilt diseases and Fo47 colonizing plants endophytically without any disease symptoms. Their distinct effects on plants, combined with their minimal genetic diversity (the two strains belong to the same species), should facilitate the identification of meaningful genotype–phenotype correlations.

In addition to being a good model system, *F. oxysporum* is of great agricultural importance, as it is listed among the top 10 most researched fungal pathogens for food production (Dean et al., 2012). Collectively, this group of filamentous fungi causes devastating vascular wilt diseases in over 100 crop species, leading to annual yield losses of billions of dollars (Ma et al., 2013). One notorious example is the recent Panama disease outbreak in banana caused by *F. oxysporum* f. sp. *cubense* Tropical Race 4 (Viljoen et al., 2020). Information accumulated over the past 10 years has provided a clear picture of compartmentalization of the *F. oxysporum* genome: A core genome component that is conserved and vertically transmitted performs essential housekeeping functions, and an accessory genome that is believed to have been initially acquired horizontally mediates unique host–fungal interactions (Yang et al., 2020; Ma et al., 2010, 2013; Zhang; Vlaardingerbroek et al., 2016b, 2016a; DeIulio et al., 2018; Hane et al., 2011; Williams et al., 2016; Galazka and Freitag, 2014; Armitage et al., 2018; van Dam et al., 2016; Dong et al., 2015).

Using an unbiased approach and taking advantage of two recently released high-quality genome assemblies of Fo47 and Fo5176 (Wang et al. 2020; Fokkens et al. 2020), we employed metatranscriptomics to dissect how Arabidopsis plants react to two *F. oxysporum* isolates with distinct lifestyles during the early course of infection. We demonstrated that endophytic infection suppresses host immunity but activates plant nutrient assimilation. By contrast, pathogenic infection activated defense response but suppressed plant developmental functions. Genome comparison of the two isolates revealed unique accessory chromosomes that harbor genes enriched for fungal virulence and detoxification in Fo5176, and cell signaling and nutrient sensing in Fo47. Our study showed that while for both plants and *F. oxysporum*, most genes displayed a similar expression pattern during infections, a small number of genes displayed transcription plasticity between endophytic and pathogenic infections, perhaps leading to the different interaction outcomes.

## RESULTS

### A pathosystem that reveals both endophytic and pathogenic interactions

To dissect beneficial versus pathogenic fungal–plant interactions, we inoculated Arabidopsis plants with two *F. oxysporum* strains, the beneficial (endophytic) strain Fo47 and the pathogenic strain Fo5176. The pathogenic fungus Fo5176, initially isolated in Australia (Thatcher et al., 2012; Chen et al., 2015), causes vascular wilt in several Brassicaceae plants, including *A. thaliana* (Thatcher et al., 2009; Ma et al., 2010). The endophytic strain Fo47 was originally isolated from disease-suppressing soils (Alabouvette, 1999) and has been used as a biocontrol agent to prevent disease from soil-borne pathogens by inducing the production of plant secondary metabolites and priming host resistance (Aimé et al., 2013; Olivain et al., 2006; Benhamou et al., 2002; Benhamou and Garand, 2001; Veloso and Díaz, 2012).

We adopted a robust and reproducible root-dipping protocol to inoculate 14-d-old Col-0 plants with a suspension of fungal spores (Thatcher et al., 2012). Plants inoculated with Fo5176 developed typical yellowing and wilting symptoms, visible at 6 d post inoculation (DPI) (Figure 1A). By then, the fungal hyphae had advanced into the stele of infected roots (Figure 1B), as revealed by staining with 5-bromo-4-chloro-3-indoxyl-α-L-arabinofuranoside (X-ARA), which is hydrolyzed to a blue precipitate by a fungal-derived enzyme (Diener, 2012). Almost all Fo5176-infected plants died within 3 weeks of inoculation (Figure 1C). By contrast, plants inoculated with Fo47 not only stayed healthy (Figure 1A, C), but also showed increased aboveground biomass (Wilcoxon rank-sum test, *p* < 0.001), when compared to mock-inoculated plants (Figure 1D), suggesting that Fo47 may have a growth-promoting effect. X-ARA staining of plant roots inoculated with Fo47 determined that fungal hyphae were restricted to the root outer layers (Figure 1B).

**Figure 1.**
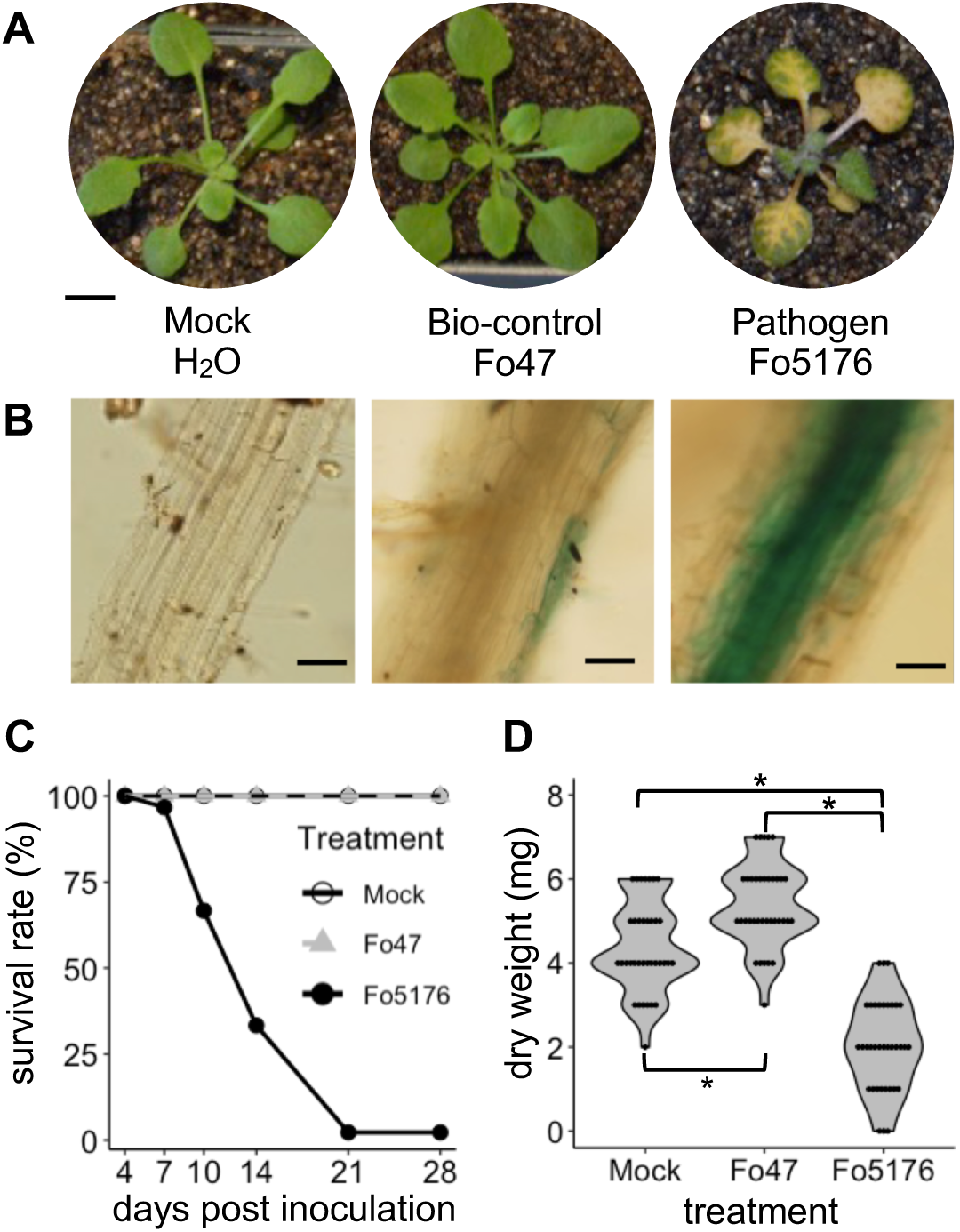
Compatible versus incompatible Arabidopsis interaction with an endophytic (Fo47) versus a pathogenic (Fo5176) *F. oxysporum* strain. **A.** *F. oxysporum* Fo5176 causes typical wilt symptoms on Arabidopsis Col-0 plants, while Fo47-infected and mock-inoculated plants with water do not exhibit any symptoms. Photographs were taken at 7 d post inoculation (DPI), and representative plants are shown. **B.** Micrographs of Arabidopsis roots mock-inoculated with water or infected with Fo47 or Fo5176 at 5 DPI and stained with X-ARA to reveal *F. oxysporum* hyphae. Scale bar represents 1 mm. **C.** Survival analysis assay illustrating the survival rates of Arabidopsis plants mock-inoculated with water or infected with Fo47 or Fo5176 at six time points, from 4 to 28 DPI. Ninety plants were assayed per treatment. **D.** Summary of shoot dry biomass of Arabidopsis plants mock-inoculated with water or infected with Fo47 or Fo5176 at 6 DPI. Statistical significance was determined by Kruskal–Wallis and Wilcoxon rank-sum tests. Asterisk indicates statistical significance at *p* < 0.001. Thirty-six plants were assayed per treatment.

By comparing to a sister species *F. verticillioides*, accessory chromosomes were identified in both the Fo47 (Wang et al., 2020a) and Fo5176 (Fokkens et al., 2020) genomes, in addition to the 11 core chromosomes (Figure 2), vertically inherited from the common ancestor shared between these two sister species 10–11 million years ago (Ma et al., 2013). The Fo47 genome had one accessory chromosome (chromosome 7, with a length of 4.25 Mb), while the Fo5176 genome had four (chromosomes 2, 14, 16, and 18) (Figure 2). The combined length of accessory chromosomes/regions in Fo5176 was 21.63 Mb, including large segments (size > 1 Mb) of chromosomes 4, 10, 11, and 13 that shared no syntenic block with the *F. verticillioides* genome. Fo47 and Fo5176 accessory chromosomes were enriched in repetitive sequences (Supplemental Figure 1), a common property observed from all accessory chromosomes (Yang et al., 2020). Fo47 accessory genes were significantly enriched for cell signaling and nutrient sensing functions, whereas Fo5176 genes were enriched for functions relating to virulence and detoxification (Supplemental Data Sets 1 and 2). As these two genomes share an almost identical core sequence, we hypothesized that distinct accessory chromosomes in each genome may play important roles in the distinct phenotypic outcomes (disease versus growth promotion).

**Figure 2.**
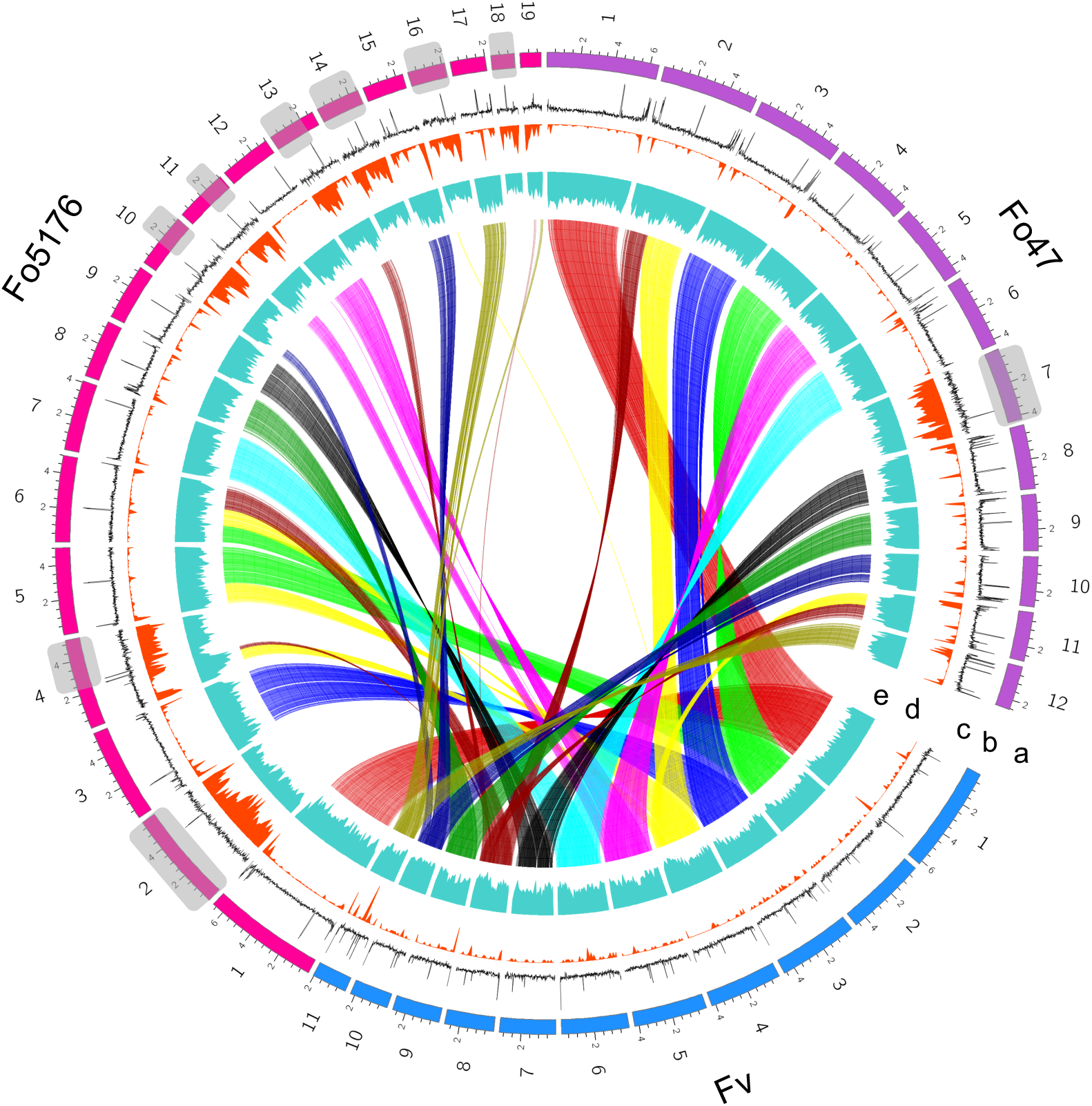
Comparative genomics reveals unique sets of accessory chromosomes in *F. oxysporum* Fo47 and Fo5176. Synteny of genome assemblies between *F. verticillioides* (*Fv*) and the two selected *F. oxysporum* strains. Track a: distribution of karyotypes of assembled chromosomes; track b, GC density; track c, density of transposable elements (TEs) calculated in 10-kb windows; track d, gene density, calculated in 100-kb windows. Track e shows syntenic blocks. Relationships are shown through linking syntenic block genes (gene number >10) in each genome pair. Core chromosomes can be identified through synteny between *Fv* and each *Fo* strains, whereas accessory chromosomes and regions show no or reduced synteny. Chromosomes 2, 14, and 18 and large segments (size > 1 Mb) of chromosomes 4, 10, 11, 13, and 16 in Fo5176 and chromosome 7 in Fo47 show no synteny with the *Fv* genome and are thus identified as accessory regions, characterized by their high TE density and low gene density.

### Reprogramming of the plant transcriptome in response to a fungal pathogen or endophyte

To examine the transcriptional regulation underlying the distinct endophytic and pathogenic interactions of the two strains (Figure 1), we sequenced the fungal and host plant transcriptomes from Arabidopsis plants inoculated with Fo47 or Fo5176. Infected plants were sampled at 12, 24, 48, and 96 h post inoculation (HPI), in parallel with plants mock-inoculated with water at 12 HPI as a control. We harvested root tissues for transcriptome deep sequencing (RNA-seq). Dual RNA-seq data were analyzed using an in-house pipeline to calculate the transcript levels of plant and fungal genes.

About half of all annotated Arabidopsis genes (16,544 out of a total of 32,833) were differentially regulated in at least one of 18 comparisons between different time points for the same interaction type (12, 24, 48, and 96 HPI; 12 comparisons), between different interaction types at the same time point (beneficial versus pathogenic; four comparisons), or between endophytic or pathogenic interactions and the mock control at 12 HPI (two comparisons). These differentially expressed genes (DEGs) revealed several interesting patterns (Figure 3A).

**Figure 3.**
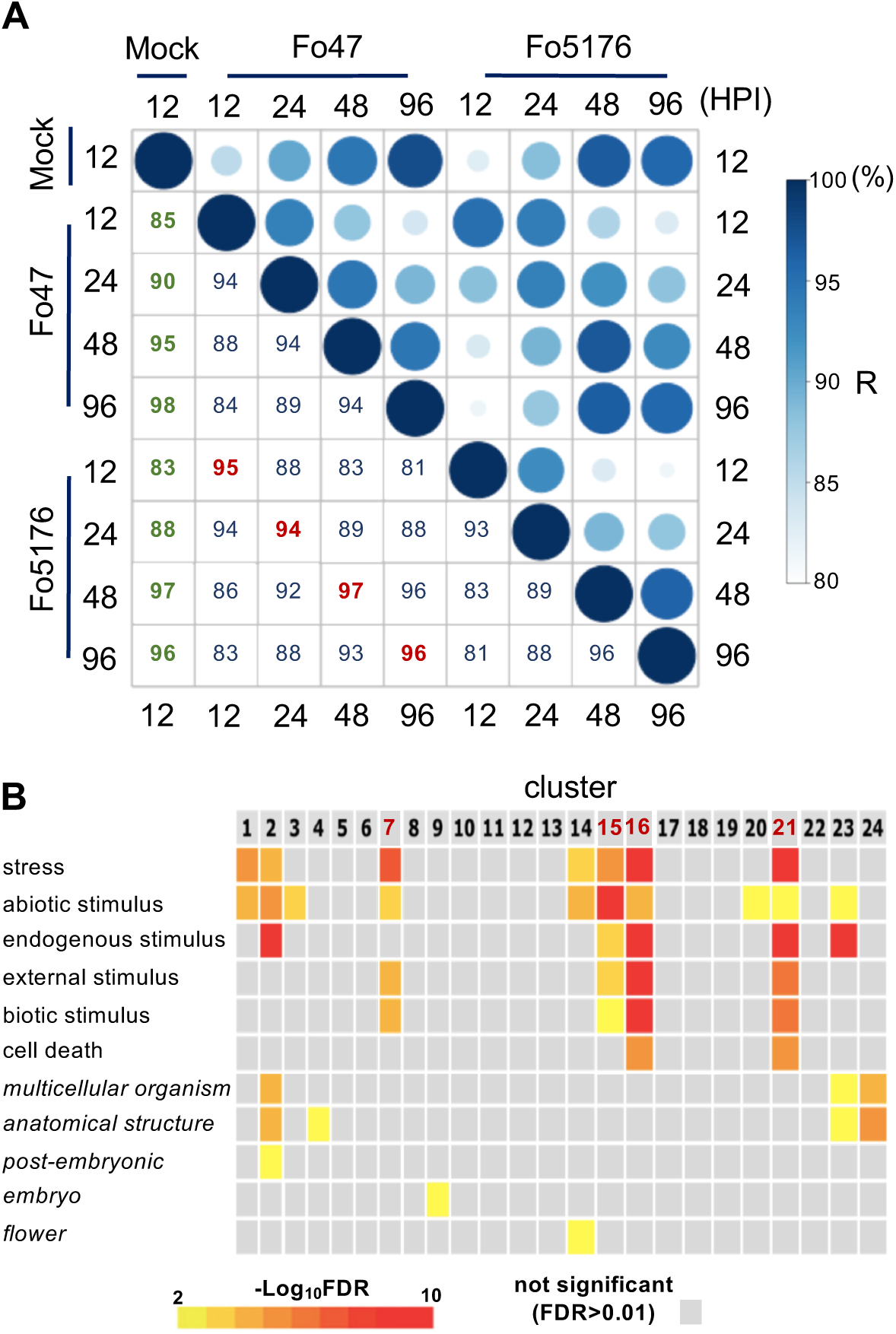
Expression profiling of Arabidopsis roots inoculated with an endophytic versus a pathogenic *F. oxysporum* strain. **A.** Extent of correlation between Arabidopsis differentially expressed genes (DEGs) in Fo47*-* and Fo5176-infected plants across the different time points at 12, 24, 48, and 96 h post inoculation (HPI). Correlation coefficients (converted to percentages) are scaled to the sizes and colors of the circles. **B.** Gene Ontology (GO) enrichment of 24 gene clusters from K-means clustering of Arabidopsis DEGs. The color scale of the heatmap represents the significance level of GO enrichment for biological processes related to stimuli response and developmental processes, expressed as –Log_10_(false discovery rate [FDR]). Four clusters, C7, C15, C16, and C21, highly enriched for stimuli responses and deprived of developmental regulation, are highlighted in red and defined as immunity clusters.

First, we observed a strong correlation between patterns of gene expression for both treatments at the same time points, despite clearly distinctive endophytic and pathogenic phenotypes (Figure 1). The Pearson’s correlation coefficients (PCCs) for the four comparisons between plants infected with either the beneficial or the pathogenic fungal strain at each time point were very high, with values of 0.95, 0.94, 0.97, and 0.96 at 12, 24, 48, and 96 HPI, respectively (labeled in red in Figure 3A), suggesting that a small subset of genes contribute to the observed phenotypic differences. Global clustering analysis using the 16,544 Arabidopsis DEGs yielded 24 co-expression gene clusters (Figure 3B, Supplemental Figure 2, Supplemental Data Set 3). A total of 10,014 genes within 12 clusters had similar expression patterns at all time points (see Supplemental Data Set 3), accounting for 60.5% of all DEGs. Considering the fact that 16,289 genes were either not expressed or not changed, we concluded that 6,544 (∼20% of all) genes held answers to the transcriptional reprogramming between these two treatments.

We also observed significant transcriptional reprogramming within each interaction over time. For samples inoculated with Fo47, PCC scores decreased from 0.94 (between 12 and 24 HPI) to 0.84 (between 12 and 96 HPI) as infection progressed. Similarly, PCC values dropped from 0.93 (between 12 and 24 HPI) to 0.81 (between 12 and 96 HPI) for Fo5176-inoculated plants. We then compared each fungal interaction pairwise at each time point to identify reciprocal DEGs (Supplemental Figure 3A), yielding 1,009, 642, 59, and 403 genes that were preferentially expressed in Fo47-infected plants and 868, 1,172, 604, and 425 plant genes in Fo5176-infected plants at 12, 24, 48, and 96 HPI, respectively (Supplemental Figure 3B). Notably, plant genes that were preferentially expressed during the endophytic interaction were enriched in Gene Ontology (GO) terms such as cell cycle, cell growth, development, response to stimuli, and cellular transport. Moreover, the genes associated with each enriched GO term showed a temporal wave as the infection course progressed, with genes involved in cell cycle highly enriched at the early stages of infection, but with a diminishing contribution that was consecutively replaced by genes related to development at around 24 HPI, response to stimuli at 48 HPI, and transport at 96 HPI (Supplemental Figure 3C). Conversely, genes preferentially induced in response to the pathogenic fungus were consistently enriched in GO terms mainly related to defense responses, with no obvious underlying temporal pattern (Supplemental Figure 3C).

Second, when compared to the mock-inoculated samples, plants inoculated with Fo47 or Fo5176 both displayed drastic transcriptional reprogramming at the earliest time point of this study (12 HPI), as these comparisons had the lowest PCCs of 0.85 for Fo47 and 0.83 for Fo5176. As time from initial inoculation progressed, however, the transcriptomes of all plants became much more similar, with PCCs rising to 0.98 for Fo47 and 0.96 for Fo5176 (labeled in green in Figure 3A). This observation indicated that the outcome of the plant–host interaction might be decided as early as 12 HPI. To begin to dissect the critical transcriptional reprogramming taking place at 12 HPI in both endophytic and pathogenic interactions, we conducted a careful analysis to identify genes that are not only differentially expressed between the two treatments, but also differentially expressed relative to mock-inoculated samples. This analysis resulted in the identification of genes that were specifically upregulated or downregulated in fungus-infected samples. These four plant gene sets consisted of 140 upregulated and 422 downregulated genes specifically in response to Fo47 infection, and 286 upregulated and 767 downregulated genes in response to Fo5176 infection.

Functional analysis of these genes using GO enrichment and network analyses (Figure 4, Supplemental Data Set 4–7) confirmed previous observations of fungal–plant interactions but also revealed unexpected findings. As expected, we observed significant suppression of genes related to plant growth by the pathogenic strain Fo5176 (Figure 4A), including genes associated with the cell cycle, cell wall organization, plant-type cell wall biosynthesis, and microtubule-based processes. Genes upregulated early in response to Fo5176 infection were highly enriched in toxin and indole metabolism, as well as small molecule biosynthesis (Figure 4B), possibly reflecting the initial upheaval brought upon by the infection. For the endophytic interaction, we noticed a significant suppression of immunity-related functions, including plant defense/immunity and jasmonic acid response (Figure 4C). This data therefore also suggested that the endophytic strain Fo47 attenuates plant defenses. Among the genes induced by the endophyte, we were pleased to see that several define a module related to nitrate metabolism and anion transport (Figure 4D), which would be consistent with the promotion of plant growth by Fo47 (Figure 1D).

**Figure 4.**
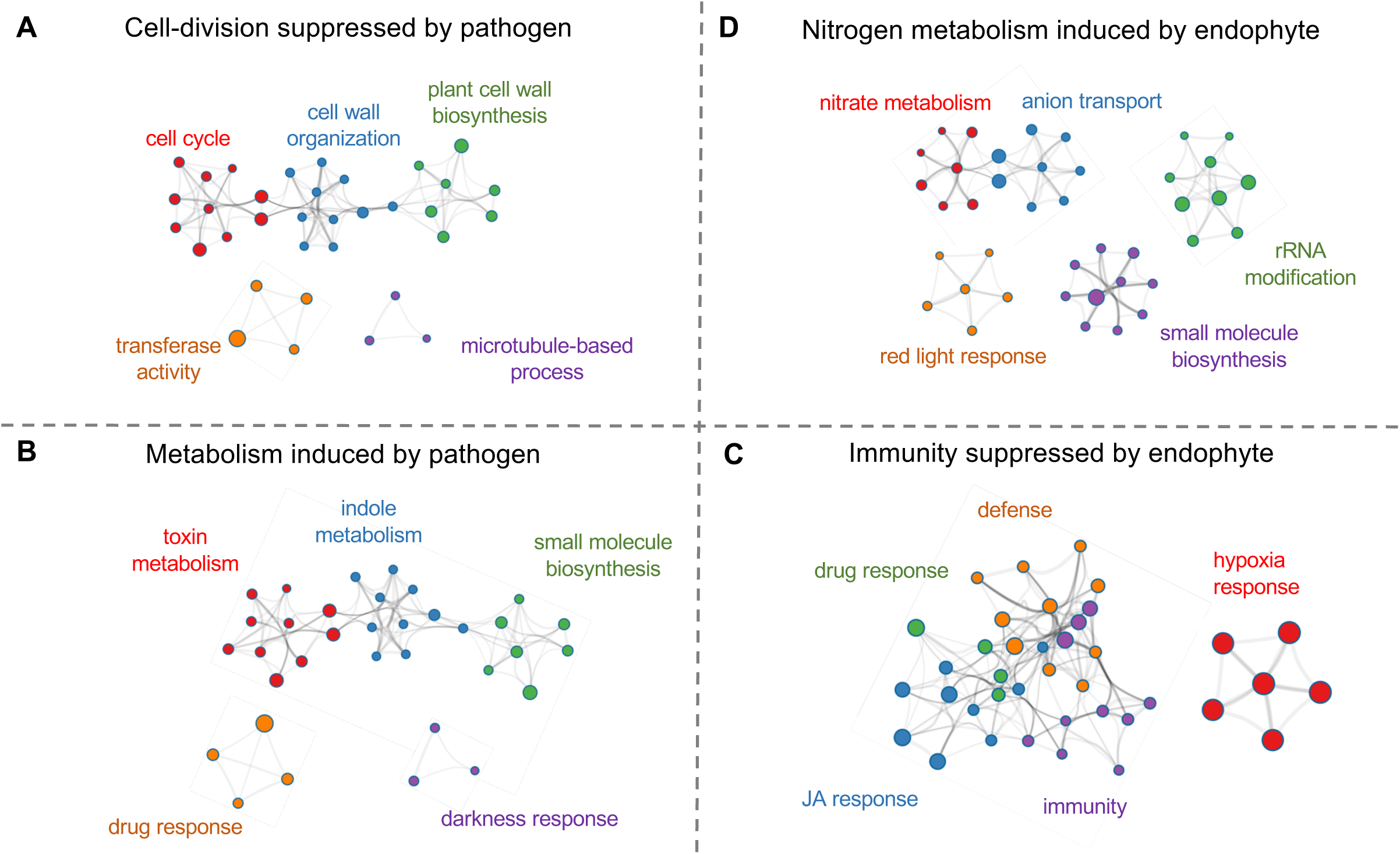
A summary of transcriptomic changes occurring at 12 HPI. GO enrichment analysis and visualization were performed on four datasets representing up- and down-regulation by Fo47 and Fo5176, respectively. Nodes represent the GO categories with enrichment, while edges exist when two GO categories share the same genes. The nodes labeled in the same color represent the GO terms that belong to a master term as labeled. The size of the nodes is scaled to the number of genes within each GO term in each figure section. **A.** Pathogen suppression: Arabidopsis genes with expression when infected by Fo47 smaller than when infected by Fo5176 and when infected by water (47 < 5176 and 47 < mock) **B.** Pathogen induction: 5176 > 47 and 5176 > mock **C.** Endophyte suppression: 47 < 5176 and 47 < mock **D.** Endophyte induction: 47 > 5176 and 47 > mock

A nitrate–CPK (Ca^2+^-sensor protein kinase)–NLP (Nin-like protein) signaling pathway was previously reported (Liu et al., 2017) that activated the expression of 394 genes and repressed another 79 genes in response to exogenous nitrate treatment. We examined whether our clusters of DEGs (Figure 3B) showed an overrepresentation of genes differentially regulated by this nitrate signaling pathway. Cluster 21 included the most downregulated genes from this pathway, with eight genes (*p*-value = 2.56e-04, two-sided Fisher’s exact test) that were downregulated in both interactions, with stronger suppression by the endophyte (Supplemental Figure 2, Supplemental Data Set 8). Of the 394 upregulated genes in the nitrate pathway, 329 were differentially expressed in our data set, with 251 assigned to clusters. Of those, over half were significantly enriched in five clusters: C5 (20 genes, *p*-value = 2.12e-05), C6 (14 genes, *p*-value = 1.07e-05), C8 (40 genes, *p*-value = 1.46e-14), C16 (16 genes, *p*-value = 7.37e-04), and C23 (38 genes, *p*-value = 6.62e-21). These 251 genes, representing a majority of the genes upregulated in the nitrate signaling pathway, were induced by both the endophyte and the pathogen but exhibited stronger responses in the context of endophytic inoculations (Supplemental Figure 2).

Notably, cluster C23 included *NLP1*, encoding a transcription factor involved in the nitrate– CPK–NLP signaling pathway (Liu et al., 2017), as well as *NITRATE TRANSPORTER2.1* and *2.2* (*NRT2.1*, *NRT2.2*), *NITRATE REDUCTASE1* (*NIA1*), and *NITRITE REDUCTASE1* (*NIR1*), all major components of the pathway that were upregulated when compared to the mock-inoculated sample (Supplemental Figure 4). Out of 24 previously reported transcription factors that control transcriptional regulation of nitrogen-associated metabolism and growth (Gaudinier et al., 2018), 16 of them were assigned to our clusters (Supplemental Data Set 9), including *WUSCHEL RELATED HOMEOBOX14* (*WOX14*) and *LOB DOMAIN-CONTAINING PROTEIN4* (*LBD4*) in cluster C23. Collectively, this analysis suggests that nitrogen signaling is involved in the *F. oxysporum*–Arabidopsis interaction and the endophyte may enhance the nitrogen signal and hence change the course of the plant response.

### Perturbation of plant immunity

To better understand how the endophyte and the pathogen perturb plant immunity via shared and distinct responses, we carefully investigated the 24 co-expression clusters based on their global patterns of expression. Four clusters, C7, C15, C16, and C21, showed enrichment (*p* < 0.05) for GO terms related to immunity and defense responses; the same clusters also lacked GO terms related to development (Figure 3C). Compared to plant PTI and ETI networks (consisting of 1,856 PTI-related and 1,843 ETI-related genes) previously constructed using a machine learning algorithm (Dong et al., 2015), three clusters (C15, C16, and C21) were enriched for both PTI and ETI genes, whereas cluster C7 was primarily enriched in PTI response genes (two-sided Fisher’s exact test *p* < 0.05) (Supplemental Data Sets 10 and 11). This suggests a transcriptional plasticity of plant immunity in responding to the endophytic and pathogenic *F. oxysporum*.

### Conserved immune response toward an endophyte and a pathogen

Cluster C15 comprised 1,290 genes and was the largest immunity-related cluster, with nearly identical plant transcriptome responses following Fo47 and Fo5176 inoculation. Indeed, genes from cluster C15 were initially strongly upregulated at 12 HPI in both interactions and gradually returned to an expression level comparable to that of mock-inoculated plants as infection progressed (Figure 5A). C15 was most significantly enriched in PTI genes (*p-*value = 2.66e-72), reflecting the general plant perception of fungal signals derived from both pathogenic and symbiotic organisms (*e.g.* MAMPs). Cluster C15 indeed included many immunity-related genes involved in fungal perception, signal transduction, and transcriptional regulation, including *ERECTA*, a RLK that regulates stomatal patterning and immunity (Sopeña-Torres et al., 2018); *RECOGNITION OF PERONOSPORA PARASITICA5* (*RPP5*), which encodes a putative NLR protein that confers resistance to *Peronospora parasitica* (Noël et al., 1999; Parker et al., 1997); *RESPONSIVE TO DEHYDRATION 21A* (*RD21A*), which encodes a cysteine proteinase with peptide ligase and protease activity that is involved in immune responses against the necrotrophic fungal pathogen *Botrytis cinerea* (Lampl et al., 2013); and *NUCLEAR FACTOR Y*, *SUBUNIT B3* (*NF-YB3*), which encodes a transcription factor activated by endoplasmic reticulum (ER) stress responsible for the regulation of stress responses (Liu and Howell, 2010).

**Figure 5.**
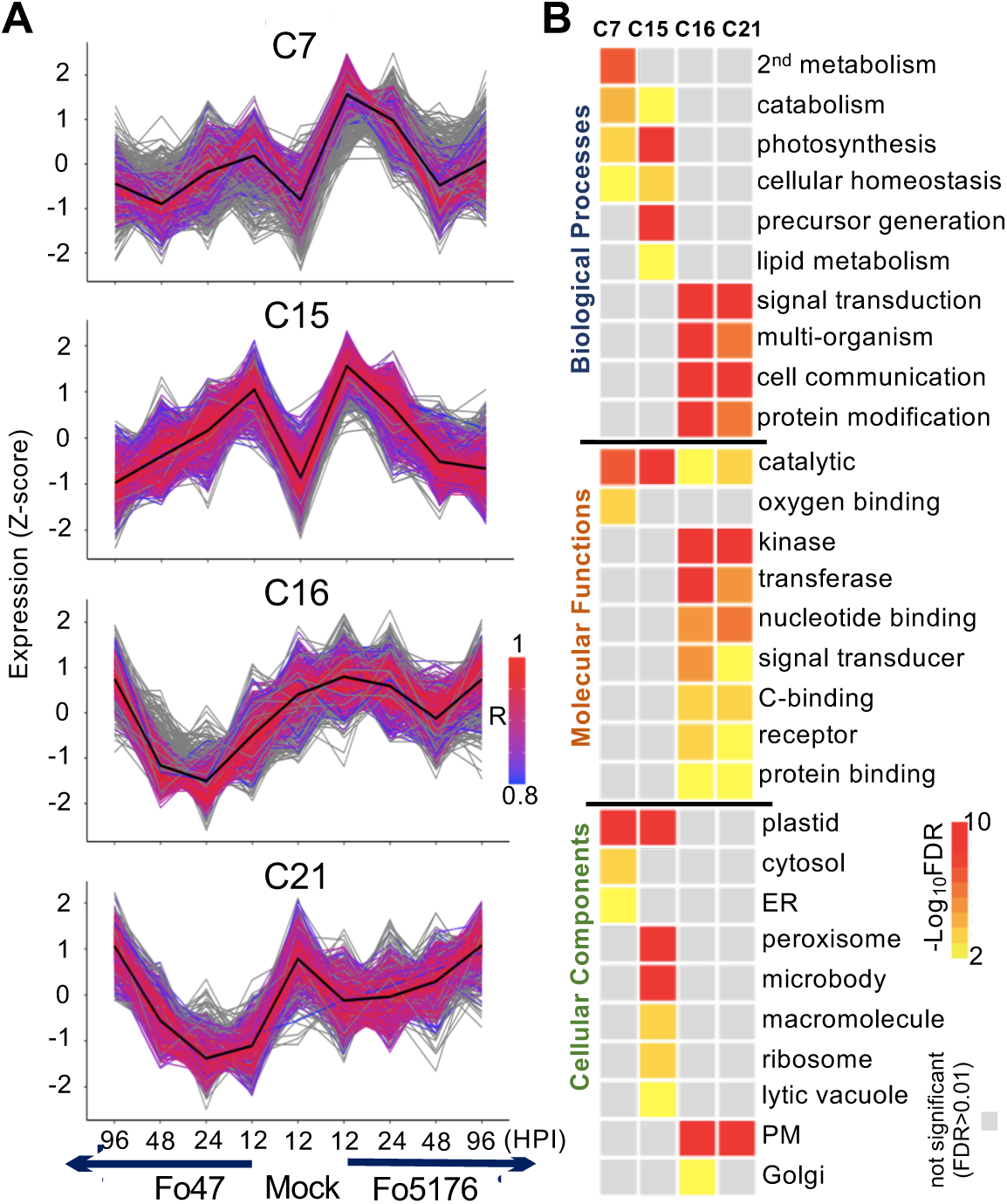
Expression and GO enrichment of immunity gene clusters. **A.** Expression profile of immunity gene clusters. Color scale indicates the correlation of expression between genes and the cluster centroids. Genes that were removed from the clusters before functional analysis due to the expression correlation with centroid lower than (or equal to) 0.8 are shown in gray. Enrichment of pathogen-associated molecular pattern-triggered immunity (PTI) and effector-triggered immunity (ETI) genes (defined by Dong et al. 2015) is indicated (in all labeled cases, *P* < 1E−7, by Fisher’s exact test). Gene number within the four clusters is as follows: C7, 422; C15, 1,290; C16, 615; C21, 766. **B.** GO enrichment analysis of immunity gene clusters for biological processes, molecular functions, and cellular components. Color scale of the heatmap represents the FDR. Stimuli responses, developmental processes (overlapping with Figure 2C), and redundant GO terms were removed.

Cluster C15 was also highly enriched in genes with functions related to the chloroplast/plastid (*p-*value = 9.2e-89, Figure 5B, Supplemental Data Set 12). An organelle essential for plant photosynthesis, chloroplasts have recently come to the forefront as key players in plant immune responses (Göhre et al., 2012; Serrano et al., 2016), possibly functioning as a signaling hub that links the initial recognition of diverse pathogens at the PM and signal transduction to the nucleus to orchestrate transcriptional reprogramming in response to infection (Medina-Puche et al., 2020; Chan et al.; de Souza et al., 2017; Wang et al., 2016; de Torres Zabala et al., 2015; Liu, 2016). It is unlikely that chloroplast-related genes from cluster C15 represent artifacts caused by the manipulation of roots in the light during harvesting, as these genes were expressed at low levels in mock-inoculated plants, although they were subjected to the same inoculation and harvesting procedure. Our observations are also consistent with a previous report in which strain Fo5176 was shown to induce the expression of Arabidopsis genes normally involved in photosynthesis in root tissues at 1 DPI (Lyons et al., 2015). Pathogens may thus interfere with host chloroplast/plastid functions to manipulate host immunity in their favor. We know very little about the possible role played by chloroplasts during endophytic colonization.

### Stronger induction of plant PTI responses by the pathogen

We hypothesized that a subset of plant immune responses against the pathogen and endophyte would differ, given their distinctive phenotypes, even though most clusters showed the same pattern during endophytic and pathogenic responses. Indeed, three immunity-related clusters, C7, C16, and C21, exhibited distinct patterns between the pathogen and the endophyte (Figure 5A). Cluster C7 (422 genes), which was primarily enriched in genes associated with PTI, exhibited a stronger induction by Fo5176 infection than by Fo47, despite being induced by both strains (Figure 5A). Several GO terms were shared between clusters C7 and C15, such as chloroplast/plastid-related functions (Figure 5B), possibly reflecting fine-tuning of the initial recognition of conserved fungal signals.

Interesting genes included *PEROXIDASE37* (*PRX37*), encoding a putative apoplastic peroxidase that generates H_2_O_2_ primarily in the vascular bundles for host defense (Pedreira et al., 2011), and *PENETRATION2* (*PEN2*), encoding an atypical tyrosinase required for broad-spectrum resistance to filamentous plant pathogens (Fuchs et al., 2016). Also included in cluster 7 were genes with dual functional roles in immunity against different pathogens. For instance, *PATATIN-LIKE PROTEIN2* (*PLP2*) promotes cell death and facilitates *Botrytis cinerea* and *Pseudomonas syringae* infection in Arabidopsis (La Camera et al., 2005), whereas it confers host resistance to Cucumber mosaic virus (Camera et al., 2009). *KUNITZ TRYPSIN INHIBITOR1* (*KTI1*), a trypsin inhibitor referred as an antagonist involved in the negative regulation of programmed cell death that mediates susceptibility in *Erwinia carotovora* but has an opposite function in *Pseudomonas syringae* pv *tomato* DC3000 (Li et al., 2008).

### Suppressed plant immunity in the presence of the endophyte

In contrast to clusters C7 and C15, both clusters C16 (615 genes) and C21 (766 genes) exhibited stronger suppression of expression by the endophyte (Figure 5A). We also observed a unique and specific suppression of plant immunity by the endophyte from the GO term enrichment and network analyses at 12 HPI described above (Figure 4C). While clusters C7 and C15 showed minimal overlap of enriched GO terms, clusters C16 and C21 shared many terms, including signal perception and transduction, protein–protein interaction, and PM localization (Figure 5B). Out of 70 genes identified as contributing to danger sensing and signaling systems (Zhou and Zhang, 2020), we detected 12 genes in cluster C16 and another 12 genes in cluster C21 (Table 1 and Supplemental Data Set 13). For instance, PRRs and downstream components in cluster C16 include *EFR*, *BAK1*, *LYK5*, and *CERK1*; and *PEPR1*, *PEPR2*, *FERONIA*, and *RBOHD* in cluster C21. NLRs and downstream signaling components in cluster C16 include *ADR1-L1/ADR1-L2* and *RPM1*, *PAD4*, and *RPS4* in cluster C21. In summary, a strong suppression of diverse immunity-related genes is unique to the endophytic interaction, suggesting that modulation of plant immunity may contribute to the different outcomes.

**Table 1.**
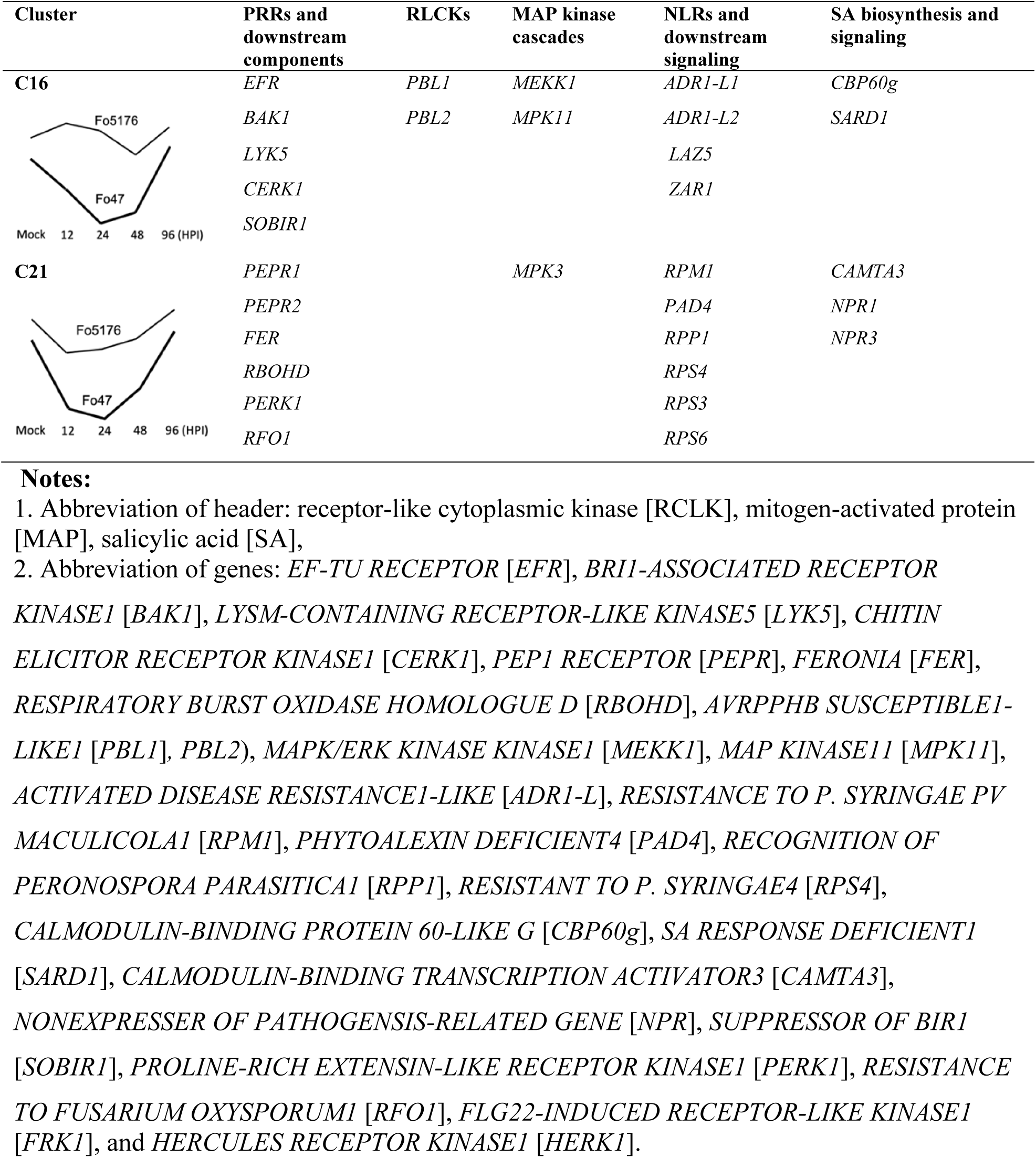
Genes that play important roles in danger sensing and signaling suppressed by the endophytic interaction

### A systematic analysis of PTI- and ETI-sensing genes induced by the endophyte and the pathogen

Overall, we observed strong host immune responses when challenged with either the endophyte or the pathogen, involving complex signal perception and signal transduction cascades. Distinct responses included the suppression of plant growth and the induction of plant defenses by the pathogenic strain Fo5176 and the attenuation of host immunity with the concomitant induction of nitrogen metabolism by the endophytic strain Fo47 (Figure 4). To further dissect the plant immunity pathways involved in these two interactions, we conducted a systematic analysis of the expression profiles of genes encoding RLPs/RLKs and NLR proteins (See supplementary data 14 and 15 and examples in Table 1).

### RLP/RLK genes

The Arabidopsis genome encodes 533 RLPs/RLKs, as determined by the MAPMAN Mercator annotation (Schwacke et al., 2019). Of those, 311 were assigned to our clusters of DEGs (Supplemental Data Set 14). In addition to the two immunity clusters C16 (37 genes, *p*-value = 4.76e-11) and C21 (30 genes, *p*-value = 2.11e-05), whose expression is repressed by the endophyte, these *RLP/RLK* genes were also enriched in clusters C11 (14 genes, *p*-value = 6.31e-04) and C12 (11 genes, *p*-value = 6.31e-04). Their expression appeared to be repressed by both the endophyte and the pathogen to varying degrees. Characterized defense-related *RLK* genes include *SUPPRESSOR OF BIR1* (*SOBIR1*) and *EFR* in cluster C16 and *RESISTANCE TO FUSARIUM OXYSPORUM1* (*RFO1*) and *PROLINE-RICH EXTENSIN-LIKE RECEPTOR KINASE1* (*PERK1*) in cluster C21. *RFO1* encodes a protein that confers a broad-spectrum resistance to *Fusarium* (Diener and Ausubel, 2005). Cluster C16 also included genes encoding LysM receptor-like kinases CERK1 and LYK5 (Cao et al., 2014), which are essential for the perception and transduction of the chitin oligosaccharide elicitor. Although not significantly enriched, 26 *RLP/RLK* genes grouped in cluster C15, including *FLG22-INDUCED RECEPTOR*-*LIKE KINASE1* (*FRK1*) and *HERCULES RECEPTOR KINASE1* (*HERK1*), which are activated in response to both pathogenic and endophytic *F. oxysporum* strains.

### *NLR* genes

Many plant NLRs are commonly identified as resistance proteins that act as surveillance molecules recognizing pathogen effectors that target the host machinery. In accordance with the so-called “guard” model, NLRs then trigger an ETI response (Cao et al., 2014). The Arabidopsis genome encodes 160 NLR proteins (Baggs et al., 2020), of which we identified 119 genes among our data set of DEGs (84 were assigned to clusters). Among these NLR genes, about 40% clustered across four immunity clusters, cluster C21 (18 genes), C16 (14 genes), C15 (11 genes), and C7 (4 genes), and were uniquely enriched in clusters 21 (*p*-value = 2.58e−07) and 16 (*p*-value = 6.82e−06), both of which include genes specifically repressed in the endophyte (Supplemental Data Set 15). The 11 NLR genes in cluster C15 included five known resistance genes (AT1G63880, AT1G61190, RPP4, RPP5, and RPP8) against oomycete and fungal pathogens (Goritschnig et al., 2012; McDowell et al., 2005; Staal et al., 2006; van der Biezen et al., 2002).

Notably, most of the *NLR* genes that were enriched in endophyte-suppressed clusters C16 and C21 are not functionally characterized. Nevertheless, characterized NLRs represented by genes in cluster C16 included two apoplast/chloroplast-localized ADP-binding immune receptors (ADR1-L1 and ADR1-L2) (Dong et al., 2016). Also belonging to cluster C16 were the two effector-induced resistance genes *LAZARUS5* (*LAZ5*) and *HOPZ-ACTIVATED RESISTANCE1* (*ZAR1*) (Barbacci et al., 2020; Baudin et al., 2017), conferring resistance to a *Pseudomonas syringae* strain expressing the AvrRPS4 and Hop effectors, respectively. *NLR* genes in cluster C21 included disease resistance proteins RPS3, RPS4, and RPS6, which provide specific resistance against *P. syringae* pv. *tomato* carrying the avirulence genes *AvrRPS3*, *AvrRPS4*, and *AvrRPS6*, respectively (Narusaka et al., 2009; Kim et al., 2009; Bisgrove et al., 1994). Repression of the expression of these *NLR* genes by the endophyte again supports the idea that the danger sensing and signaling systems underlying the responses to Fo47 and Fo5176 are distinct.

### Accessory chromosomes in two strains harbor genes induced during infection and with distinct biological functions

The distinct plant responses at both the phenotypic and transcriptome levels, resulting from inoculation with the two *F. oxysporum* isolates, is in no doubt related to genomic differences between the two strains. Even though both strains belong to the same species complex and share an ∼46-Mb core genome, each strain also carries distinct accessory chromosomes (Figure 2). The Fo47 accessory chromosome 7 harbored 1,299 predicted genes (7.2% of total predicted genes) (Supplemental Data Set 16), 757 of which were expressed and 160 were strongly induced (false discovery rate (FDR) < 0.05) during one or more time points of the infection.

To explore the function of these accessory genes encoded in the endophytic strain Fo47, we next analyzed the functional domains they encoded. After excluding genes that encoded proteins with transposase-like domains or unknown domains, we highlighted five enriched PFAM domains: regulator of G-protein signaling domain (PF00615), nitric oxide (NO)-binding membrane sensor involved in signal transduction (PF03707), basic Leucine Zipper (bZIP) transcription factor (PF00170), chromodomain (PF00385), and bromodomain (PF00439) (Figure 6A, Supplementary Figure 5A). The enrichment of functional domains involved in cell signaling and the apparent lack of enrichment for domains related to virulence suggested that the Fo47 accessory chromosome contains genes with functions that are well suited to a nonpathogenic life style. Transcriptome analysis showed that nine Fo47 genes encoding proteins with the bacterial signaling protein domain (PF03707) are most highly induced at 24 and 48 HPI. Domain PF03707 plays a role in sensing oxygen, carbon monoxide (CO), and NO (Galperin et al., 2001). The Fo47 genome has the highest number of genes encoding a PF03707 domain, with nine genes, compared to other filamentous fungi such as *Aspergillus nidulans* (1), *Neurospora crassa* (1), *Magnaporthe grisea* (1), *F. graminearum* (2), *F. verticillioides* (2), and *F. solani* (3) (Galagan et al., 2005; Cuomo et al., 2007; Dean et al., 2005, Ma et al., 2010), as well as other FOSC members (3 ∼ 7, average 4) (DeIulio et al., 2018). Notably, six of the nine genes reside on accessory chromosome 7, making it a major contributor to the expansion of this gene family within this strain (Supplemental Figure 6).

**Figure 6.**
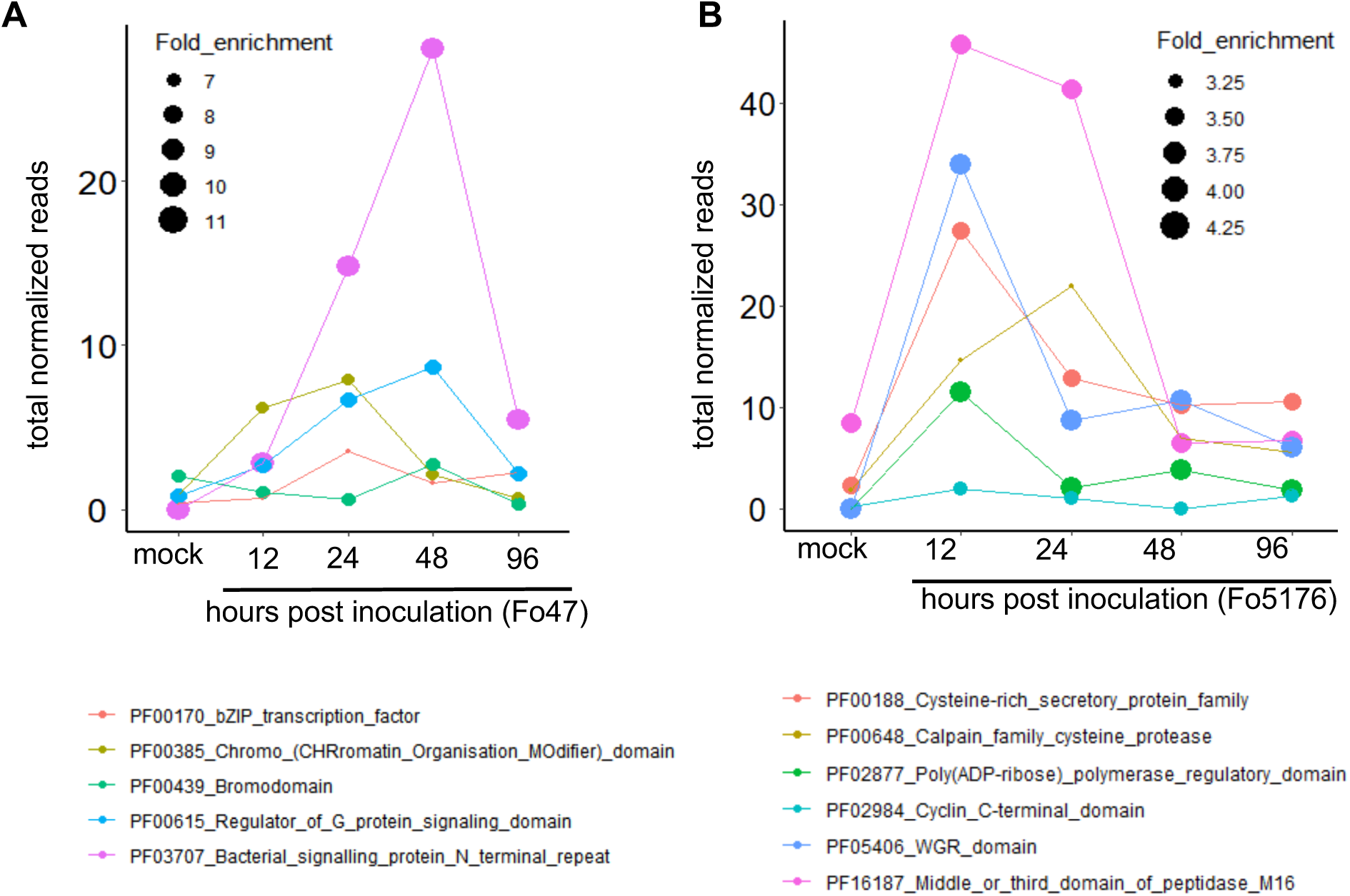
Distinct biological functions for induced AC genes in the endophyte Fo47 and the pathogen Fo5176. Fold enrichment refers to the ratio of the proportion of genes on the accessory chromosomes (ACs) with a specific term over the proportion of genes in the whole genome with a particular term (adjusted *p*-value <0.05). **A.** Significantly induced Fo47 accessory genes are represented in five enriched PFAM domains, including regulator of G-protein signaling domain (PF00615), NO-binding membrane sensor involved in signal transduction (PF03707), bZIP transcription factor (PF00170), chromo domain (PF00385), and bromodomain (PF00439) containing proteins. **B.** In Fo5176, six PFAM domains are significantly enriched and induced at different stages of infection course: cysteine-rich secretory protein family (PF00188), Calpain family cysteine protease (PF00648), peptidase M16 (PF16187), poly (ADP-ribose) polymerase regulatory domain (PF02877), cyclin C-terminal domain (PF02984), and WGR domain (PF05406).

By contrast, accessory chromosomes and regions from the pathogenic strain Fo5176 contributed 4,136 predicted genes (23% of total predicted genes) (Supplemental Data Set 17), of which 3,502 were expressed and 1,140 were strongly induced during one or more time points of the infection. Genes located in accessory regions in Fo5176 encoded proteins that were enriched for 42 PFAM domain terms. We noticed six PFAM domains that were highly enriched at different stages of the infection course and whose encoding genes were highly expressed: cysteine-rich secretory protein family (PF00188), Calpain family cysteine protease (PF00648), peptidase M16 (PF16187), poly (ADP-ribose) polymerase regulatory domain (PF02877), cyclin C-terminal domain (PF02984), and WGR domain (PF05406) (Figure 6B, Supplementary Figure 5B). Most of these domains are likely associated with microbial pathogenesis or detoxification; their associated genes were induced during infection (Figure 6B). In particular, members of the cysteine-rich secretory protein (CAP) superfamily (PF00188) have a wide range of biological activities, including fungal virulence, cellular defense, and immune evasion (Schneiter and Di Pietro, 2013). For example, the *F. oxysporum* CAP family protein Fpr1 is a PR-1-like protein that is important for the virulence of strain Fol4287 (Prados-Rosales et al., 2012). The Fo5176 genome encodes 15 CAP family members, significantly more than Fo47 and other comparable fungal species (average: 5 members) (Supplemental Figure 7). Phylogenetic analysis of CAP proteins showed that four CAP members formed a core group shared by Fo5176 and Fo47. However, a separate clade of six CAP family proteins was expanded in Fo5176 and encoded by Fo5176 accessory chromosomes (Supplemental Figure 7). These results highlight the distinctive functions of, and roles played by, accessory chromosomes in the nonpathogenic strain Fo47 and the pathogenic strain Fo5176. These differences might provide the mechanistic basis that allows Fo47 to specialize in host sensing and benefit its host as an endophyte, while the pathogenic Fo5176 specializes in host invasion and killing.

## DISCUSSION

We performed a comparative study of infection by an endophytic (Fo47) and a pathogenic (Fo5176) strain of *F. oxysporum* in the context of the *F. oxysporum* Arabidopsis system, which revealed the transcriptional plasticity of plant defense responses. The pathosystem we developed combines the extensive knowledge of plant immunity in Arabidopsis and one of the most damaging fungal pathogens for agriculture, *F. oxysporum*. Strain-specific interactions with a common host are likely dictated by the accessory chromosomes from each *F. oxysporum* genome, which allows a comparative study that minimizes genetic differences between strains to address the underlying mechanism that results in distinct phenotypes (growth promotion or disease or even death). Up to 50% of crop losses in the USA can be attributed to soil-borne pathogens (Raaijmakers et al., 2009), and our results provide a foundation for the development of technologies to enhance plant health, sustain a healthy ecosystem, and feed a continuously growing human population.

We employed comparative metatranscriptomics over the course of early infection to systematically capture temporal transcriptional changes in both the host and the interacting microbes. Our results may be summarized along four main axes, as illustrated in Figure 7.

**Figure 7.**
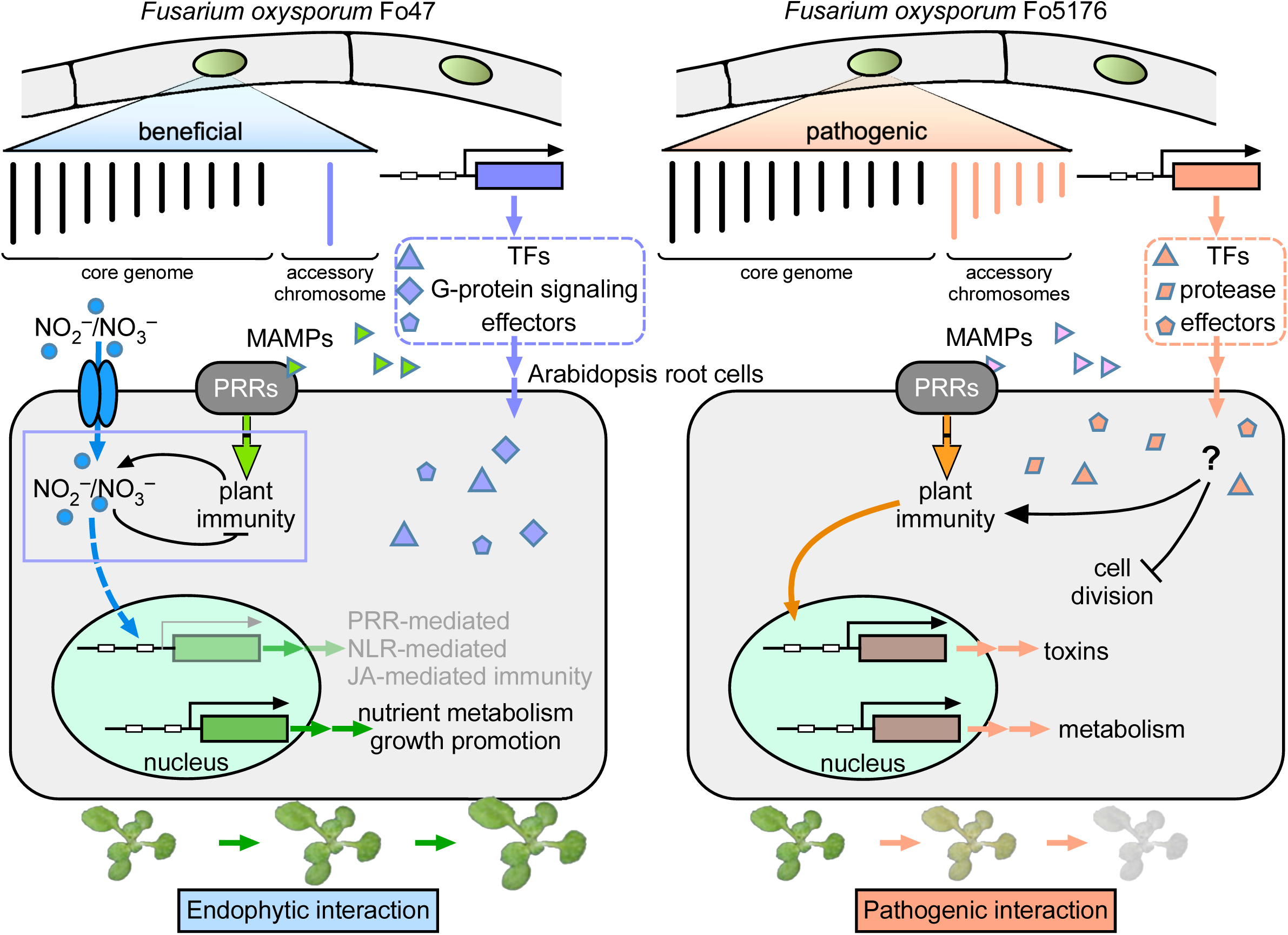
Model of transcriptomic plasticity in beneficial and antagonistic plant–fungal interactions. Molecular response of *Arabidopsis thaliana* plants challenged with an endophyte Fo47 and a pathogen Fo5176, two *Fusarium oxysporum* strains share a core genome of about 46 Mb, in addition to their unique accessory chromosomes. Distinct responding genes depict the transcriptional plasticity, as the pathogenic interaction activates plant stress responses and suppresses plant growth/development-related functions, while the endophyte attenuates host immunity but activates plant nitrogen assimilation. The differences in reprogramming of the plant transcriptome are linked to accessory genes encoded by the two closely related fungal genomes.

First, host transcriptional responses were strikingly similar at all time points, regardless of the obvious phenotypic differences seen after infection of Arabidopsis plants by Fo47 and Fo5176. Of all clusters of DEGs, cluster C15 exhibited a strong and early induction of genes, followed by a return to an expression level comparable to that of control samples. This cluster captured 26 Arabidopsis *RLP/RLK* genes, as well as 269 PTI and 159 ETI response genes, suggesting that both strains initially elicit a similar MAMP response, which would not be surprising as they belong to the same species.

Second, our data revealed rapid transcriptional reprogramming at the beginning of the interactions. While the majority of plant genes exhibited a common expression pattern during the two infections, a small subset of plant genes displayed divergent gene expression profiles. By far the most striking difference was observed at 12 HPI, when the GO biological processes for genes uniquely induced by Fo47 or Fo5176 reflected almost opposite responses. The endophytic strain Fo47 stimulated nitrogen metabolism and suppressed host immunity, whereas the pathogenic strain Fo5176 stimulated host immune responses and toxin metabolism, but repressed functions related to plant growth and development. We propose that this distinct expression profile, reflected in the early divergence of the host transcriptome, is the result of plasticity of the host transcriptome when facing an endophyte or a pathogen. We hypothesize that the perception by the host of distinct fungal signals occurs shortly after inoculation and is followed by the rapid activation of downstream signaling cascades. Our results also stress the importance and necessity of sampling early during the establishment of a fungal–host interaction to better capture the full extent of the underlying temporal dynamics.

Third, Fo47 inoculation resulted in suppression of genes related to plant defense and induced genes related to plant growth, in agreement with the trade-off between growth and defense. It has been reported that plants can channel nitrogen resources towards production of defense-related compounds when confronted with pathogens (Ullmann-Zeunert et al., 2013). For instance, allele polymorphism at the single locus *ACCELERATED CELL DEATH6* (*ACD6*) can dictate distinct difference between growth and defense among different Arabidopsis ecotypes (Todesco et al., 2010). Further characterizing the *F. oxysporum*–Arabidopsis pathosystem should illuminate the mechanism(s) by which nutrients are allocated in relation to plant defense.

Finally, we observed an agreement between plant infection phenotypes and distinctive gene functions associated with fungal accessory chromosomes. While upregulated fungal accessory genes were primarily enriched in proteins with roles in cell signaling and nutrient transport in the endophyte Fo47, they were enriched for virulence and detoxification in the pathogen Fo5176, likely contributing to the contrasting phenotypes of plants infected by these two *F. oxysporum* strains.

In conclusion, time-resolved comparative metatranscriptomics can be used to characterize transcription regulation when the model plant Arabidopsis is challenged with an endophyte and a pathogen of the same fungal species. We showed both the conservation and plasticity of the plant and fungal transcriptomes and how they may relate to the distinctive genomic features associated with each fungal genome. The Arabidopsis and *F. oxysporum* pathosystem developed here is likely to become an ideal system to characterize plant recognition and response mechanisms against soil-borne root fungi. We believe this system will be pivotal in enriching our understanding of the molecular mechanisms necessary to enhance vascular wilt resistance not only in Arabidopsis, but also in crops that are under threat by *F. oxysporum* pathogens.

## METHODS

### Plant and fungal growth

*Fusarium oxysporum* strains Fo5176 and Fo47 were routinely cultured on potato dextrose agar (BD, New Jersey USA) at 28°C under a 12-h-light/12-h-dark photoperiod. Fungal spores were collected from 5-d-old cultures in potato dextrose broth (BD, New Jersey USA) by passing the liquid culture through a double layer of sterile cheesecloth, followed by centrifugation of the flow-through at 3,000 *g* for 15 min at room temperature. Fungal spores were mixed with an appropriate volume of sterile deionized water to prepare the spore suspension (concentration: 1 × 10^6^ spores/mL) for infection assays.

### Plant infection assay

Seeds of Arabidopsis (*Arabidopsis thaliana*) accession Columbia-0 (Col-0) were obtained from the Arabidopsis Biological Resource Center (ARBC, Ohio State University) and were surface sterilized in 1 mL of 70% (v/v) ethanol three times, 5 min each, followed by one wash with 50% (v/v) bleach for 5 min. After removing the bleach solution, seeds were rinsed with 1 mL of sterile distilled and deionized water, and stratified for 3–4 d in darkness at 4°C. Seeds were planted into pots filled with an autoclaved mixture of fine-grain play sand: MetroMix 360:vermiculite in a 1:2:1 ratio, watered with distilled deionized water, and covered with a clear plastic lid to retain a high humidity for 3 d in the growth chamber with the following settings: 24°C, 14 h light/10 h dark, and a light intensity (T8 fluorescent and incandescent bulbs) ranging from 89 to 94 μmol·m^−2^·s^−1^. After 3 d, the plastic lid was removed, and seedlings were allowed to grow for 11 additional days prior to inoculation with *F. oxysporum* microconidia. Plants were 14 d old at the time of inoculation and had at least four fully expanded true leaves. For *F. oxysporum* infection, the roots of 14-d-old Arabidopsis plants were dipped for 30 s in a 1 × 10^6^ fungal spores/mL suspension of Fo5176 or Fo47, or in sterile dH_2_O for the mock control. Inoculated plants were planted in autoclaved potting mix and moved to a growth chamber set to 28°C with the same photoperiod as above.

### RNA preparation, sequencing, and data analysis

Roots from infected plants at 12, 24, 48, and 96 h post inoculation (HPI) were harvested from five plants per treatment and time point for total RNA isolation. For control samples, roots from the same number of control plants were collected at 12 HPI. Fungal cultures from Fo5176 and Fo47 fungal mycelia were harvested after 5 d from liquid cultures for RNA extraction. Three biological replicates were produced for each treatment. Total RNA was extracted using the ZR Soil/Fecal RNA Microprep Kit (Zymo Research, CA, Cat. R2040) following the manufacturer’s protocol, and the RNA quantity and quality were assessed using a NanoDrop 2000 and Agilent 2100 Bioanalyzer. Illumina TruSeq stranded mRNA libraries were prepared and sequenced on an Illumina HiSeq2000 platform at the Broad Institute (Cambridge, MA). One replicate each for infected plant samples inoculated with Fo47 at 12 HPI and Fo5176 at 24 HPI failed, as did one replicate for Fo47 and Fo5176 mycelia samples; these four conditions are therefore only represented by two replicates and were used for downstream processing and analysis.

Paired-end RNA-seq reads were first assessed for quality by FastQC 0.10.1 (Andrews, 2010). RNA-seq data were analyzed using the HISAT, StringTie, and DESeq2 pipelines (Pertea et al., 2016; Love et al., 2014). Briefly, reads were mapped to reference genomes of Arabidopsis (annotation version Araport11, (Cheng et al., 2017)), Fo5176 (Like Fokkens et al., 2020), and Fo47 (Wang et al., 2020a) using HISAT2/2.0.5 (Kim et al., 2015). Mapped reads were used to quantify the transcriptome by stringTie/1.3.4 (Pertea et al., 2015). Read count normalization and differential gene expression analysis were conducted using DESeq2/1.27.32 with a maximum FDR of 0.05 (Love et al., 2014). Corrplot/0.84 was used to visualize the correlation in gene expression profiles between different conditions. Read counts of differentially expressed genes (DEGs) were first averaged per condition and then normalized by log transformation as log_2_(normalized read count + 1), and then correlations were calculated. Clustering analysis on per-condition averaged, log-transformed, and Z-scaled read counts was performed using the K-means clustering algorithm ‘Lloyd’ (R function K-means) and then visualized in ggplot2/3.3.0.

### Functional analysis and visualization

GO enrichment analysis (plant GO slim) of Arabidopsis gene clusters and reciprocal DEG analysis were conducted with the singular enrichment analysis (SEA) tool of agriGO v2 (Du et al., 2010; Tian et al., 2017) using the Arabidopsis TAIR10 annotation. We applied a hypergeometric test, combined with Hochberg (FDR) multi-test adjustment method to discover enriched GO terms at a significance level of 0.01 with a minimum of three mapping entries. Comparisons of different enrichment results were performed using cross-comparison of SEA (SEACOMPARE). We generated PFAM annotations for *F. oxysporum* 5176 and 47 by InterproScan, following a standard annotation pipeline (Jones et al., 2014). PFAM enrichment in proteins encoded by fungal genes was performed in TBtools (Chen et al., 2020) using Fisher’s exact test with FDR < 0.05. We performed a custom analysis in Metascape (Zhou et al., 2019), with the options minimum overlap of 3, *p*-value cutoff of 0.01, and minimum enrichment of 1.5 for the discovery of GO term enrichment and network visualization of Arabidopsis DEGs at 12 HPI. The top five terms (with the smallest *p*-values) were selected, and the terms that shared the gene entries (forming edges) were visualized. The visualization was further polished in Cytoscape/3.8.0 (Shannon, 2003).

### Synteny and phylogenomic analysis

Synteny was detected by Basic Local Alignment Search Tool for nucleotides (BLASTN), with parameters above 50 kb coverage and 98.5% sequence identity, and visualized as a Circos plot (Krzywinski et al., 2009). *OrthoFinder* was used to identify orthologous pairs across compared genomes and to construct a genome-based phylogenetic tree. The divergence times between species were estimated using the PL method with r8s (Taylor and Berbee, 2006). CAFE (Computational Analysis of gene Family Evolution) v.3 (Han et al., 2013) was used to test whether protein family sizes were compatible with a stochastic birth and death model, and the Viterbi algorithm in the CAFE program was used to assign *p*-values to the expansions/contractions experienced at each branch and using a cutoff of *p* < 0.05.

### Accession Numbers

RNA-seq data generated in this study were deposited in the NCBI Short Read Archive (SRA) with accession number GSE87352.

## Supplemental Data

**Supplemental Data Set 1.** PFAM domain enrichment of genes located on accessory chromosomes in Fo47.

**Supplemental Data Set 2.** PFAM domain enrichment of genes located on accessory chromosomes in Fo5176.

**Supplemental Data Set 3.** Statistics and gene list of 24 co-expression gene clusters.

**Supplemental Data Set 4.** Host genes that are preferentially upregulated by Fo47 at 12 HPI.

**Supplemental Data Set 5.** Host genes that are preferentially downregulated by Fo47 at 12 HPI.

**Supplemental Data Set 6.** Host genes that are preferentially upregulated by Fo5176 at 12 HPI.

**Supplemental Data Set 7.** Host genes that are preferentially upregulated by Fo47 at 12 HPI.

**Supplemental Data Set 8.** Cluster assignment and annotation of nitrate response genes that were differentially expressed in our study.

**Supplemental Data Set 9.** Cluster assignment and annotation of transcription factors that control transcriptional regulation of nitrogen-associated metabolism and growth.

**Supplemental Data Set 10.** Overlap of PTI-responsive genes with immunity clusters.

**Supplemental Data Set 11.** Overlap of ETI-responsive genes with immunity clusters.

**Supplemental Data Set 12.** Genes within C15 that are annotated as encoding plastid-localized proteins.

**Supplemental Data Set 13.** Cluster assignment of the genes that are involved in immune signaling.

**Supplemental Data Set 14.** Cluster assignment and annotation of *RLP/RLK* genes that were differentially expressed in our study.

**Supplemental Data Set 15.** Cluster assignment and annotation of *NLR* genes that were differentially expressed in our study.

**Supplemental Data Set 16.** The 1,229 genes located on Fo47 accessory chromosomes and their expression at five selected stages.

**Supplemental Data Set 17.** The 4,136 genes located on Fo5176 accessory chromosomes and their expression at five selected stages.

## ACKNOWLEDGEMENTS

The authors thank Dr. John Manners and Donald Gardiner of CSIRO for providing the strain Fo5176; Chris Joyner, the Superintendent of the College of Natural Sciences Greenhouse at University of Massachusetts, for his helps with all experiments conducted in the greenhouse; and Patrice Patrice Salomé for preparing the summary figure.

This project was supported by Natural Science Foundation of (IOS-165241), the National Research Initiative Competitive Grants Program Grant no. 2008-35604-18800 and MASR-2009-04374 and MAS00496 from the USDA National Institute of Food and Agriculture. Data were analyzed at the Massachusetts Green High Performance Computing Center (MGHPCC). LG is also supported by a China Postdoctoral Foundation Grant (2017 M623188), the National Natural Science Foundation of China (31701739) and the Fundamental Research Fund of Xi’an Jiaotong University (1191329155). L.-J.M. is also supported by an Investigator Award in Infectious Diseases and Pathogenesis by the Burroughs Wellcome Fund BWF-1014893, and the National Eye Institute of the National Institutes of Health under award number: R01EY030150. HLY is also supported by Lotta M. Crabtree Fellowship. The funding bodies played no role in the design of the study and collection, analysis, and interpretation of data and in writing of the manuscript.

## AUTHOR CONTRIBUTIONS

Project design and oversight: LG, LJM; Providing fungal strains: CS, VE, HCK; Conducting experiments: LG, LZ, GD, AB, KV; data analysis: LG, HLY, HY, BW, LJM; Results interpretation: LG, HLY, LJM; Manuscript writing: LG, HLY, HCK, LJM; Manuscript revision: all authors; Provide funding: LG, LJM. All authors read and approved the manuscript.

## Supplemental Data

**Supplemental Figure 1. Comparison of transposable element numbers in core regions and accessory regions of Fo47 (A) and Fo5176 (B).**

Bin size = 100 kb. t-tests were performed, and *p*-values are indicated.

**Supplemental Figure 2. Co-expression gene clusters.**

Results of K-means clustering analysis, yielding 24 co-expression gene clusters. Color scale indicates the correlation of expression between genes and the cluster centroids. Genes that were removed from the clusters before further functional analysis due to the expression correlation with centroid lower than (or equal to) 0.8 are shown in gray. X axis, from left to right in each plot, dictate 96 HPI, 48 HPI, 24 HPI, 12 HPI (Fo47), Mock, 12 HPI, 24 HPI, 48 HPI, and 96 HPI (Fo5176).

**Supplemental Figure 3. Reciprocal DEGs between endophyte- or pathogen-infected Arabidopsis plants at each time point.**

**A.** Diagram of the overall analysis.

**B.** Reciprocal Arabidopsis DEGs in endophytic or pathogenic *F. oxysporum* infections at FDR < 0.05.

**C.** Gene Ontology analysis (plant GO slim terms) of biological processes and molecular functions for preferentially expressed genes in Arabidopsis plants infected with Fo47 or Fo5176 at each time point. Color scale of the heatmap represents the FDR.

**Supplemental Figure 4. Expression pattern of key components involved in nitrogen metabolism.**

Key components of the nitrogen metabolism pathway were extracted from the KEGG database; the corresponding gene expression profile is shown as a heatmap.

**Supplemental Figure 5. Expression profiles of *in planta-*induced lineage-specific genes for Fo47 (A) and Fo5176 (B), encoding carrying the PFAM domains enriched on fungal accessory chromosomes.**

**Supplemental Figure 6. Comparison of the PF03707 domain (regulator of G-protein signaling domain) between Fo47 and Fo5176, and other filamentous fungi**

**A.** Number of proteins with the PF03707 domain and number of genes encoding this domain.

**B.** Phylogenetic tree of proteins with the PF03707 domain in Fo47 and Fo5176.

**Supplemental Figure 7. Comparison of the PF00188 domain (cysteine-rich secretory protein (CAP) superfamily) between Fo47 and Fo5176, and other filamentous fungi**

**A.** Number of proteins with the PF00188 domain and number of genes encoding this domain.

**B.** Phylogenetic tree of proteins with the PF00188 domain in Fo47 and Fo5176.

**Supplementary Figure 8. Single-copy orthologs of Fo47 and Fo5176 display conservation of gene expression.**

There were 11,896 orthologs (1:1) between Fo47 and Fo5176. The Pearson correlation coefficient of Fo47 and Fo5176 ortholog expression is 0.73 (mycelia), 0.82 (12 HPI), 0.72 (24 HPI), 0.88 (48 HPI), and 0.92 (96 HPI).

